# scFates: a scalable python package for advanced pseudotime and bifurcation analysis from single cell data

**DOI:** 10.1101/2022.07.09.498657

**Authors:** Louis Faure, Ruslan Soldatov, Peter V. Kharchenko, Igor Adameyko

**Affiliations:** Department of Neuroimmunology, Center for Brain Research, Medical University Vienna, 1090 Vienna, Austria; Department of Biomedical Informatics, Harvard Medical School, Boston, MA 02115, USA

## Abstract

scFates provides an extensive toolset for analysis of dynamic trajectories comprising tree learning, feature association testing, branch differential expression and with a focus on cell biasing and fate splits at the level of bifurcations. It is meant to be fully integrated into scanpy ecosystem for seamless analysis of trajectories from single cell data of various modalities (e.g. RNA, ATAC).

**Availability and implementation:** scFates is released as open-source software under the BSD 3-Clause “New” License and is available from the Python Package Index at https://pypi.org/project/scFates/. The source code is available on Github at https://github.com/LouisFaure/scFates/

**Supplementary information:** A supplementary document is provided with a complete explanation of the underlying statistics, and two figures showing examples of analysis.

---

Pseudotime analysis is a concept firstly introduced in the early days of single cell analysis for microarray data (Magwene et al., 2003). Derived from the idea that the temporal structure of gene expression can be retrieved by looking at its geometry (Rifkin and Kim, 2002), pseudotime analysis consists in ordering cells in a single dimensional value meant to recapitulate the underlying transcriptional transition. The advent of more sensitive scRNAseq methods was accompanied by the development of two main classes of high-resolution trajectory analysis tools. One relies on probabilistic mapping of the cells, with tools such as Palantir (Setty et al., 2019), and CellRank (Lange et al., 2022). Probabilistic approaches have the advantages of being flexible and efficient in finding terminal states, with the possibility for manual selection of these. However, the results do not lead to an easily interpretable tree as each cell is given a probability value for each fate, without clearly defined branches or segments. The other class of tools relies on principal graph learning, with the most used tool being Monocle3 (Cao et al., 2019). Principal graph learning allows to reconstruct a differentiation tree with discrete segments and bifurcations, enabling easier interpretation. However, Monocle3 learns the principal graph on UMAP embedding, a low dimensional representation highly sensitive to parameters and which can represent a highly distorted view of the data (Chari et al., 2021). Moreover, none of the trajectory inference python tools provide functions dedicated to test features for association with pseudotime as well as differential expression between branches. Finally, none of the R or python packages perform statistical analysis of bifurcation point, including the discovery of early cell fate biasing factors (Bargaje et al., 2017).

Here we introduce scFates, a python package to streamline the whole process of pseudotime analysis, with flexible tree learning options, advanced feature extraction tasks and specific functions focused on bifurcation analysis. scFates was initially a based on crestree R package, developed for the analysis of cell fate dynamics in development (Soldatov et al., 2019; Krivanek et al., 2020; Faure et al., 2020). While the initial R version included a tree inference approach inspired by SimplePPT (Mao et al., 2015). scFates additionally implements ElPiGraph, another method for principal graph learning (Albergante et al., 2020), allowing investigators to impose topological constrains on trajectories to fit single curves or circular trajectories. scFates is fully compatible with scanpy ecosystem (Wolf et al., 2018) by using the anndata format, and provides GPU and multicore accelerated functions for faster and more scalable inference. scFates is divided into three main parts: (1) trajectory learning and graph operations, (2) feature significance over pseudotime testing and clustering, and (3) bifurcation analysis. All three parts implement publication-grade plotting capabilities to visualize the results.

The trajectory learning can be performed on processed single cell data at different stages of analysis, such as on normalised transcript count matrix, PCA or diffusion maps. The user has the choice of running either ElPiGraph or SimplePPT algorithms to learn a principal graph. To leverage the advantages of probabilistic mapping, scFates can also learn a principal graph on CellRank output by considering the combination of the absorption probabilities of each fate and calculated pseudotime as a manifold reduction, from which can be applied SimplePPT (fig. 1A). The latter is relevant in cases where CellRank is more successful in capturing terminal fates than principal graph methods. The resulting trajectory, composed of connected principal points, can be projected onto any low-dimensional space, such as UMAP or ForceAtlas2. To orient a reconstructed trajectory, the user can choose one or two roots manually or according to the value of a feature, such as level of expression of a known marker. Pseudotime positions of individual cells are then determined by projecting cells according to their position relative to the principal points. Thanks to the soft assignment matrix properties from SimplePPT algorithm, where each cell is assigned a probability to each principal point, scFates can generate several pseudotime mappings to account for variability. In addition, scFates provides convenient functions for selecting specific portions of the tree, by selecting starting and endpoints, or by using pseudotime. The association of any of the molecular features measured in the dataset (e.g. gene expression, ATAC peaks) with pseudotime can be tested using provided generalized additive models (GAM). For trees, such association tests are performed in a branch-specific manner. Associated features can be fitted and smoothed over pseudotime using the same model and can then be clustered using a distance metric of choice (fig. 1D). scFates also incorporates a test for bridges or transitions captured by the trajectory, based on deviations from linearity (fig. 1B). By checking whether the expression dynamics observed in the bridge can be explained by a linear mixture of the flanking populations, such test can be used to assure that the bridge is not an artifact, and uncover molecular transient features associated with these transitions (Kameneva et al., 2021). Inspired by a recent dedicated R package (Hou et al., 2021), scFates provides a function specifically designed for covariate testing is proposed to test for differential gene expression between covariates sample conditions on the same trajectory path. This test includes an amplitude difference part, as well as trend differences (fig. 1C).

**Figure 1:**
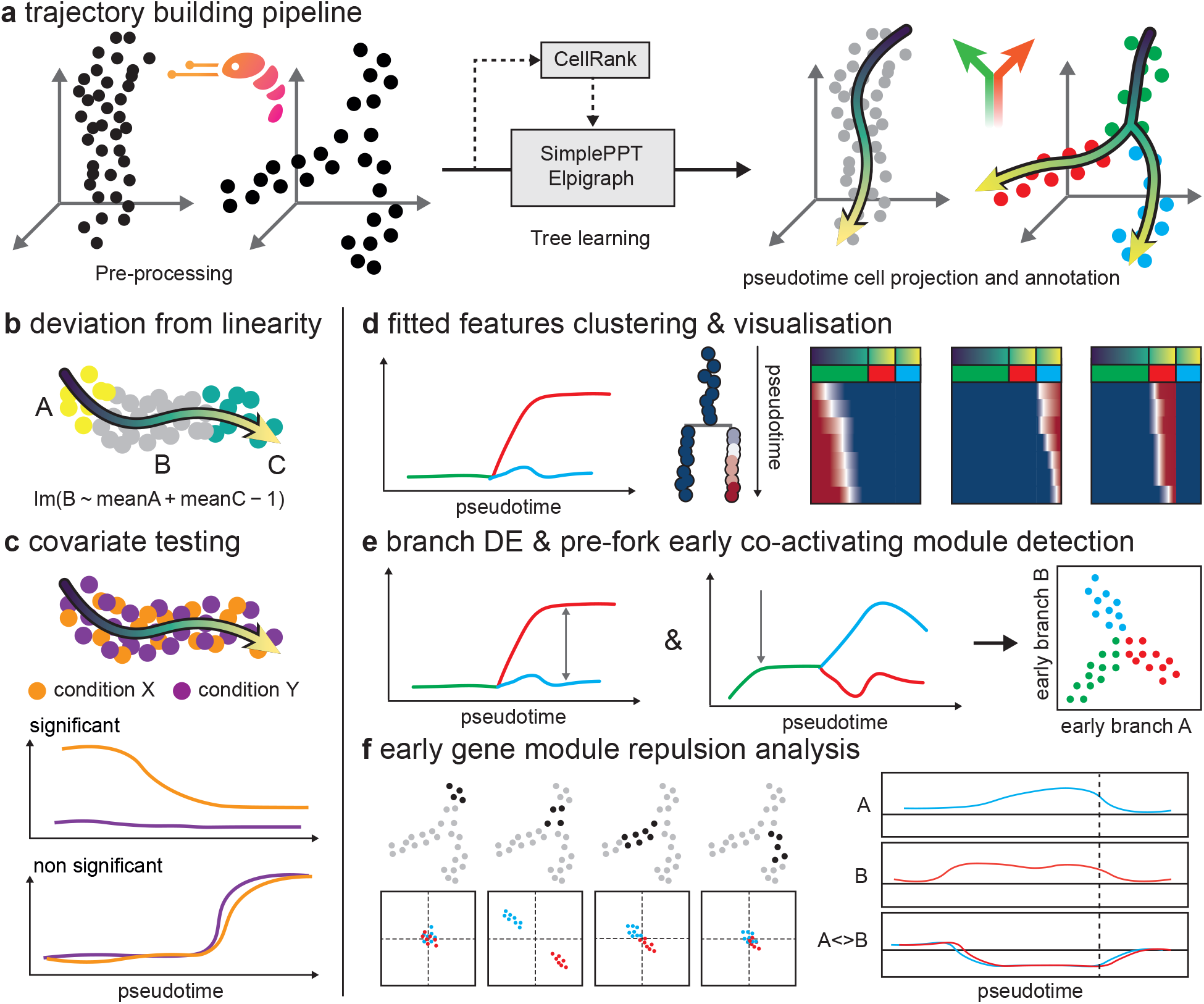
Overview of scFates functionalities: **(A)** Dataset is first pre-processed via commonly used tools such as scanpy, to generate a reduced space where the tree will be learned using either ElPiGraph, SimplePPT or a combination of CellRank absorption probabilities and SimplePPT. Pseudotime and segment assignment is then calculated for each cell. **(B)** Transitions can be tested for deviation from linearity, to assess whether the bridge is likely to be an artifact and reveal transient features. **(C)** In trajectories containing two or more conditions, changes in amplitudes and/or trends can be statistically tested for each feature. **(D)** Fitted features can be clustered and visualised in different ways. **(E)** Identification of early gene modules **(F)** Local gene to gene correlation analysis of bifurcations, including detection of intra- and inter-early modules and their repulsion.

Finally, scFates implements a set of functions for detailed analysis of bifurcations. First, early and late gene modules separating within each branch are detected by testing for differential expression between branches, and determining the timing of their deviation relative to the bifurcation (fig. 1E). scFates then identifies features of early modules that contribute to cell fate biasing prior to the bifurcation point through backtracking of their correlations in the progenitor branch (fig. 1F).

In conclusion, scFates is a fast and versatile tool for in-depth tree learning, pseudotime analysis, and characterization of bifurcation dynamics from single cell data. It can be widely applied in developmental biology, disease trajectory or perturbation analysis. Moreover, scFates can work with any type of highly dimensional data, allowing to use with modalities other than gene expression, such as single-neuron activity over time.

## Notes

https://scfates.readthedocs.io/en/latest/

https://pypi.org/project/scFates/

### Competing Interest Statement

The authors have declared no competing interest.

