## Supplementary materials for "scFates: a scalable python package for advanced pseudotime and bifurcation analysis from single cell data"

### Reconstruction of principal tree with SimplePPT

SimplePPT is available as a separate python package named simpleppt. It consists in fitting a principal tree in M-dimensional space to a set of data points $x_1, ..., x_N$. The tree is composed of principal points $z_1, ..., z_K$ arranged and connected in the same space. Precise description of the underlying mathematics is available in the related paper [1]. The output of the algorithm are three matrices (fig. 1): F, describing the position of the principal points in the same space as the datapoints (dimensions: K×M). B, an adjacency matrix defining the principal tree (dimensions: K×K). R, a soft assignment matrix linking the observed data-points to the principal points (dimensions: K×N).

Briefly, the algorithm starts with the initialization of a matrix F. These initial positions can be manually given or randomly sampled by choosing K cells as being the initial principal points. F is then fixed, and R and B matrices are solved. B is solved by obtaining the minimum spanning tree from principal point pairwise distances. R is obtained by solving the following constrained optimization problem:

$$\min_{R \in B_r} \sum_{i=1}^{N} \sum_{k=1}^{K} r_{ik}(\|x_i + z_k\|^2 + \sigma r_{ik} \log r_i k) \quad (1)$$

Which is in other words consisting in computing pairwise distance matrix P between principal points and data points. P is transformed following a similar step applied in diffusion maps algorithm [2]:

$$R = \exp{(-\frac{P}{\sigma})} \quad (2)$$

The resulting transformation is then row-normalized, leading to a matrix with each data point $i$ being assigned to a principal point k, with the following characteristics:

$$0 > r_{ik} > 1 \quad \text{and} \quad \sum\nolimits_{k=1}^{K} r_{ik} = 1 \quad (3)$$

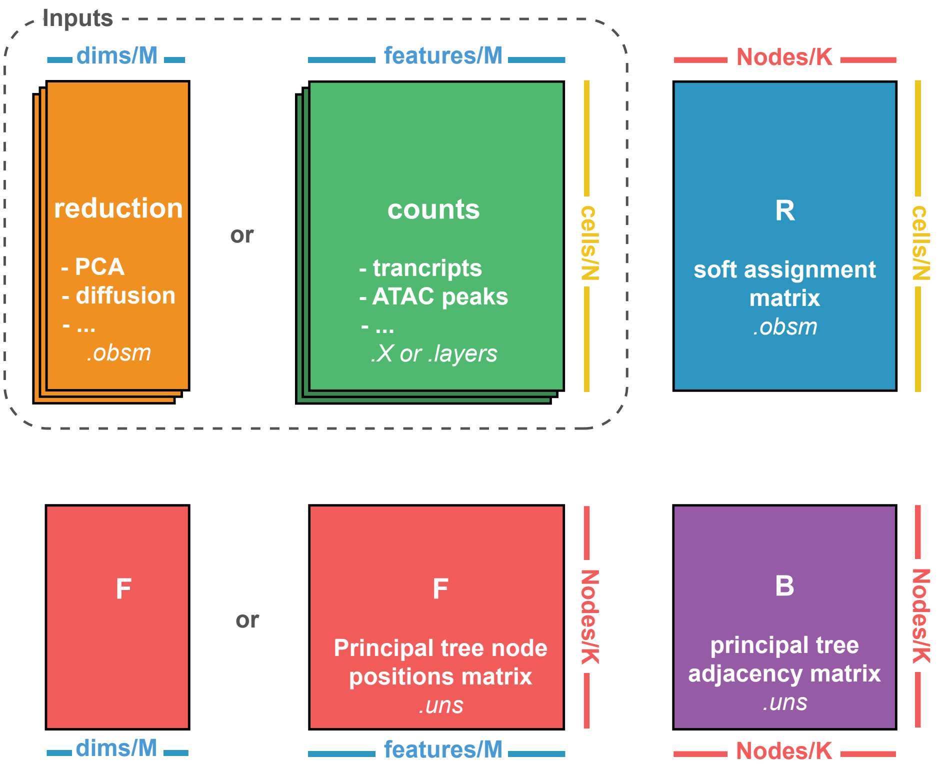


Figure 1: Schematic of the data inputs (in dashed lines) that scFates uses for tree learning. Outputs are shown out of the dashed lines. Text in italic shows the data location in anndata format

$\sigma$ in equation (2) is the sigma parameter, which acts as a gaussian kernel to control the locality of the assignments between principal points and data points. For example, a sigma too high will collapse all the principal points into the "middle" of all data points.

Once R and B are calculated, these are fixed to calculate F, by solving the following:

$$F = XR(\lambda L + \Lambda)^{-1} \tag{4}$$

Where L is the Laplacian matrix, calculated by subtracting the diagonal of the degree matrix of B by the matrix B. $\Lambda$ is the diagonal matrix of the row-sums of R. $\lambda$ is the parameter lambda which constrains the total length of the tree.

These two main steps are repeated until a defined number of steps or convergence, the latter being defined as the difference between the current F and previous F matrices is below a defined threshold. SimplePPT algorithm can be used when calling the following function:

```
scFates.tl.tree(adata, Nodes, method="ppt", ppt_sigma, ppt_lambda)
```

### Reconstruction of principal tree with ElPiGraph

ElPiGraph relies on elastic principal graphs to learn a tree in reduced space. Elastic principal graphs are based on minimization of the energy potential containing three parts:

$$U = MSE + \lambda U_E + \mu U_R \tag{5}$$

$MSE$ is the mean squared error of data approximation, $U_E$ is the sum of squared edge lengths, $U_R$ is a term minimizing the deviation of the principal graph embedment from harmonicity (generalization of linearity). $\lambda$, $\mu$ are coefficients of regularization. The structure of the graph is generated by application of a sequence of graph transformations, using operations from predefined graph grammar. The simplest graph grammar "Bisect an edge", "Add a node to a node" leads to construction of a principal tree. A more detailed explanation of the algorithm can be found in the related publication [3]. The output of the algorithm is a principal tree, curve or circle, composed of nodes and a list of edges connecting them together.

To make ElPiGraph output compatible with scFates, the node edge list is converted into the B adjacency matrix, the nodes positions are used as F matrix, and R is generated by assigning cells to their closest principal point, generating a hard assignment matrix, with the following characteristics:

$$r_{ik} \in \{0, 1\} \quad \text{and} \quad \sum_{k=1}^{K} r_{ik} = 1 \tag{6}$$

ElPiGraph algorithm can be used when calling the following functions:

```
scFates.tl.tree(adata, Nodes, method="epg", epg_lambda, epg_mu)
scFates.tl.curve(adata, Nodes, epg_lambda, epg_mu)
scFates.tl.circle(adata, Nodes, epg_lambda, epg_mu)
```

### Reconstruction of principal tree from CellRank reduction with SimplePPT

scFates can construct a principal tree on CellRank output to allow better interpretation of bifurcations. To do so, fate absorption probabilities are combined into a 2D simplex using circular posteriori projection [4]. The calculated CellRank pseudotime is concatenated into the 2D simplex to generate a 3D space, on which SimplePPT will be run. Convertion from CellRank output can be performed by calling the following function:

```
scFates.tl.cellrank_to_tree(adata, time, Nodes)
```

### Mapping of cells on the tree

Assigning a pseudotime value to cells requires first to assign that value to principal points. To do so, a root node is selected, either in a manual manner, or according to the value of a measurement (expression of marker, or other cell-wise observation). Pseudotime of a given node is calculated as the distance on the principal tree from the root to that point, in the distance metric used for tree learning. The root is selected with the following function:

```
scFates.tl.root(adata,root)
```

To map a cell onto a given tree, the first closest principal point $pp1$ is selected by using the maximum assignment value from R. The closest principal point to $pp1$, $pp2$, is also selected. The cell is then projected onto the edge connecting $pp1$ and $pp2$, by calculating a weighted average of both principal points distances to the root, using the probabilities given by R. This weighted average is considered as the pseudotime value of the cell (fig. 2 A & B.).

To perform probabilistic mapping of the cells onto the tree, we can take advantage of the fact that the soft assignment matrix R represents natural probabilistic measure as seen in (3) to assign cells onto principal points. This approach is not possible if the tree has been calulcated using ElPiGraph algorithm, due to the hard assignment matrix it generates (6). The first principal point $pp1$ is sampled according to the probabilities given by R, then the same process is performed as described in the previous paragraph.

Cell mapping assigns to each cell a pseudotime value, a segment and a milestone. The latter, inspired by dynverse, highlight the forks and tips. Cell mapping is performed by the following function:

```
scFates.tl.pseudotime(adata)
```

### Testing for features associated with the tree

Feature testing is carried out using generalized additive models (GAM) [5]. Feature expression is modeled as a function of pseudotime in a branch-specific manner, using cubic spline regression $exp_i \sim t_i$ for each branch independently. This tree-dependent model is then compared with the unconstrained model $exp_i \sim 1$ using F-test. P-values are then corrected for multiple testing, a feature is considered significant by applying a threshold on FDR values (fig. 2 C.). This association test is performed by the following function:

```
scFates.tl.test_association(adata)
```

### Covariate analysis

Features from curved trajectories or segments from trees containing cells from two or more conditions (e.g. perturbation, disease sample) can be tested for differential expression. First, for each covariates, features are tested for significant change along the pseudotime as explained in the previous section using the following function:

```
scFates.tl.test_association_covariate(adata, group_key)
```

The union of significant feature from each test are then tested for amplitude using the following GAM model:

$$g_i \sim s(pseudotime) + s(pseudotime) : Covariate + Covariate \tag{7}$$

Where $s(.)$ denotes the penalized regression spline function and $s(pseudotime) : Covariate$ denotes interaction between the smoothed pseudotime and covariate terms. From this interaction term, p-values can be extracted and then corrected for multiple testing. Features can also be tested for trend differences, comparing the model described in (7) to the following reduced one:

$$g_i \sim s(pseudotime) + Covariate \tag{8}$$

Comparison between the two model is performed with ANOVA and p-values are corrected for multiple testing. The amplitude and trend tests can be performed using the following function:

```
scFates.tl.test_covariate(adata)
```

### Linearity deviation test

To find features that specifically characterize the transitions (B) but not the cell population before (A) or after (C) the transitions, a test is performed to check whether feature expression in the transition (B) could be explained by a linear mixture of expressions of that gene in populations A and C. The feature expression profile of each cell in B was modeled as a linear combination of mean gene expression profiles in A and C with the lm function in R without the intercept term:

$$lm(B \sim meanA + meanC - 1) \tag{9}$$

For each gene in each cell in bridge B, the magnitude of the residuals not explained by the model is calculated. Mean residuals across all cells in bridge B are calculated and normalized to the standard deviation of the expression of a given gene. The obtained normalized mean residual values can be used to prioritize the genes with distinctive expression patterns in the bridge population. The test can be run with the following function:

```
scFates.tl.linearity_deviation(adata)
```

**Bifurcation analysis**

Branch specific features are first detected via amplitude testing (tl.test_fork) using the following GAM model:

$$g_i \sim s(pseudotime) + s(pseudotime) : Branch + Branch \quad (10)$$

From $s(pseudotime) : Branch$ interaction term, p-values are extracted and then corrected for multiple testing. Then, each significant feature is tested for its upregulation along the path from progenitor to derivative branch, using the linear model $g_i \sim pseudotime$, This first steps are performed using the following function:

```
scFates.tl.test_fork(adata,root_milestone,milestones)
```

Differentially expressed features are then assigned between two post-bifurcation branches with defined differences in expression and FDR cutoffs (fig. 2 D.). This is done via:

```
scFates.tl.branch_specific(adata,root_milestone,milestones,effect)
```

Finally, DE genes can be separated between early and late genes module, according to their pseudotime activation (fig. 2 G.). A gene is considered early if its pseudotime activation is before the pseudotime of the bifurcation. There are two ways to estimate the pseudotime of activation of the DE features. The first approach estimates it by separating the cells into n pseudotime bins (10 bins by default), and by calculating the relative expression rate at a specific bin:

$$r(b_t) = \frac{f(b_{t+1}) - f(b_{t-1})}{\max f - \min f} \quad (11)$$

Where $f(b)$ is the mean fitted expression at a specific bin, if the rate is higher than a defined threshold, the gene is considered to activate at the pseudotime value of the related bin. This approach can be used by calling:

```
scFates.tl.activation(adata,root_milestone,milestones)
```

The second approach uses the linear model $g_i \sim pseudotime$ in the progenitor branch only to estimate if the feature displays an upward trend before the fork. While this method does not output an estimated activation pseudotime, it is a robust alternative to separate early and late genes in more challenging dataset. This approach can be used by calling:

```
scFates.tl.activation_lm(adata,root_milestone,milestones)
```

**Local compositional heterogeneity analysis**

To study local heterogeneity along a pseudotime trajectory and near a bifurcation, cell composition around a given pseudotime $t$ is approximated by $n$ cells closest to $t$. To account for cell assignment uncertainty, the probability of a cell being close to a pseudotime point t can also estimated as a fraction of cases the cell is close to $t$ across a number of sampled cell pseudotime mappings. The related function is:

```
scFates.tl.slide_cells(adata,root_milestone,milestones,win)
```

Local gene-gene correlation reflects coordination of genes around a given pseudotime $t$, and is defined as a weighted gene-gene Pearson correlation with cells weighted according to the probability of being close to the pseudotime $t$. Local correlation of a gene $g$ with a module, is assessed as a mean local correlation of that gene with the other genes comprising the module. Similarly, intra-module and inter-module correlations are taken as the mean local gene-gene correlations of all possible gene pairs inside one module, or between the two modules, respectively. The repulsive score, reflecting antagonistic expression of competing modules at a given developmental stage, is defined as the difference between inter- and intra- module correlations. If genes inside the module correlate much stronger than they do with the competing module, such repulsive score will be close to -1, otherwise it will be closer to 0 (fig. 2 H.). Such calculation can be performed using the following function:

```
scFates.tl.slide_cors(adata,root_milestone,milestones)
```

Such function is also provided with a sliding window of overlapping cells over pseudotime, allowing to extract trends of intra- and inter- module correlations over the trajectory (fig. 2 I.). To ensure that the observed correlations do not arise due to the induction/repression trends of the considered genes, a control model can be employed, where the expression levels of each gene are permuted in windows of $n_{perm}$ cells along the pseudotime axis. The related function for such analysis are:

```
scFates.tl.synchro_path(adata,root_milestone,milestones)
```

**Formation of fate-specific modules**

In order to study decision-making process prior to the tree-reconstructed bifurcation point, a framework is provided to identify the timing of gene inclusion into its module. The approach is based on two assumptions. First, the lineage bias at a given developmental stage is manifested as coordinated expression of a subset of fate-specific genes. Second, if a gene is involved in a coordinated expression with a subset of fate-specific genes at a given developmental stage, then it will continue to be involved at later stages. In other words, the genes are progressively included into a coordinately expressed fate-specific module during differentiation, and are never excluded from the module once they join it. With these assumptions, one can infer the time of gene inclusion into a coordinated fate-specific module using a step-wise indicator function $I_g(t)$ for each fate-specific gene $g$:

$$I_g(t) = \begin{cases} 0 & \text{if } t < \text{ time of gene inclusion} \\ 1 & \text{if } t \geq \text{ time of gene inclusion} \end{cases} \tag{12}$$

This approach assesses the correlation of a gene with the time-dependent module at different time points, and identifies an inclusion time of a gene as a time at which such correlation becomes significant. Given some inclusion function $I_g(t)$, the correlation of a gene $g$ at time point $t$ with time-dependent module $M_t = \{g'|I_{g'}(t) = 1\}$ is:

$$cor(g, M_t|t) = \frac{\sum_{g' \in M_t} cor(g, g'|t)}{|M_t|} \tag{13}$$

where $g'$ is a gene from the fate-specific module $M_t$. $cor(g, M_t|t)$ is approximated by the Pearson correlation among a defined number of cells closest to time point $t$. To assess the statistical significance of $cor(g, M_t|t)$ expression of each gene can be locally permuted among a defined number of cells from which an empirical p-value can be extracted. As a result, a time-series of p-values $P_g(t)$ is generated for each gene. The time of inclusion is identified as a change-point switching from insignificant to significant p-values. More specifically, a new step function $I_g(t)$ that maximizes probability is used:

$$L = P(P_g(t_1), ..., P_g(t_n)|I_g(.)) = \prod_i P\left(P_g(t_i)|I_g(t_i)\right) \tag{14}$$

Where $P_g(t_i|1) = \frac{\alpha}{1-e^{-\alpha}} e^{-\alpha P_g(t_i}$, $P_g(t_i|0) = 1$, and $\alpha$ is a parameter that regulates the stringency of the desired significance level. The log likelihood can be rewritten as:

$$\log L = \sum_{t_i \geq \text{time of inclusion}} \left[ -\alpha P_g(t_i) + \log \frac{\alpha}{1 - e^{-\alpha}} \right] \tag{15}$$

Iterative optimization is used to identify functions $\{I_g(.)\}$, initializing them as $I_g(.) = 1$. At each step, given $\{I_g(.)\}$, for each gene $g$ we calculated $cor(g, M_t|t)$, then estimated $P_g(t)$, and then updated $I_g(t)$. In practice, the method converges in less than 10 iterations. All steps described in that section are performed by the following single function (fig. 2 J.):

```
scFates.tl.module_inclusion(adata,root_milestone,milestones)
```

**Data reproducibility**

The whole figure 2 can be fully reproduced by following the related notebook in the documentation: `https://scfates.readthedocs.io/en/latest/Tree_Analysis_Bone_marrow_fates.html`

[5] Trevor Hastie and Robert Tibshirani. *Generalized additive models*. Wiley Online Library, 1990.

[6] Manu Setty et al. "Characterization of cell fate probabilities in single-cell data with Palantir". In: *Nature Biotechnology* 37 (4 2019), pp. 451–460.

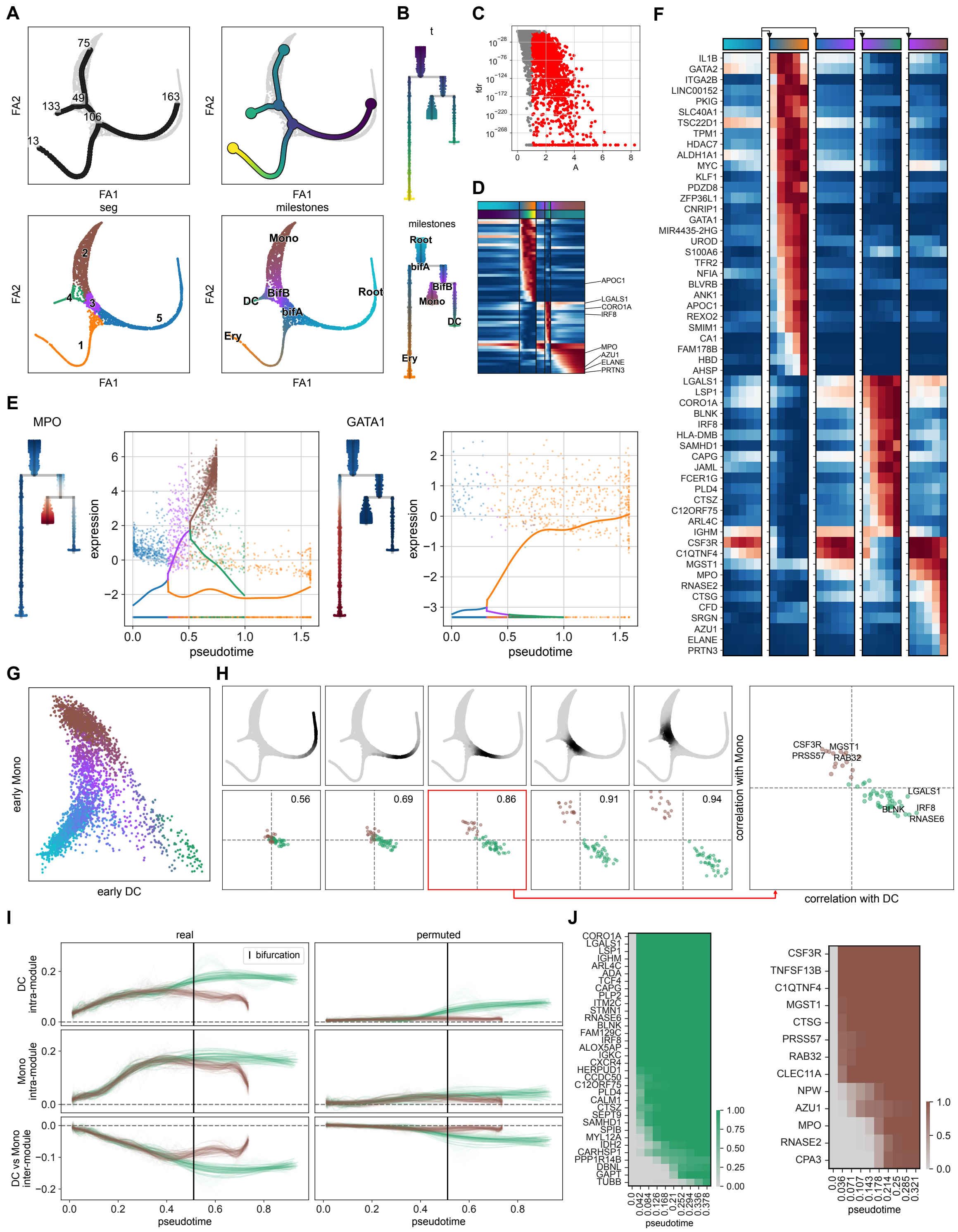


Figure 2: scFates analysis plot examples on bone marrow dataset used in Palantir study [6]. All panels have been assembled without any modification from python output: **A)** Representation of the tree before (top left) and after (top right) root selection and pseudotime calculation. Cells are assigned a segment (lower left) and a milestone (lower right). **B)** Dendrogram representation of the tree. **C)** Detection of significantly associated genes with the the tree. **D)** Top DE branch specific marker expressions showed as heatmap of cells aligned with pseudotime. **E)** Single trend plot from two known markers. **F)** Same data as shown in D, this time as a matrix plot for summarized trends over bins of pseudotime, per segment. **G)** Early gene module mean expression over cells for a bifurcating trajectory. **H)** Correlation and repulsion analysis of two competing early gene module along non-intersecting windows of cells. **I)** intra- and inter-module correlation analysis over sliding window of cells. **J)** module inclusion analysis.
